# Novel insights into the regulation of chemerin expression: role of acute-phase cytokines and DNA methylation

**DOI:** 10.1101/765834

**Authors:** Kamila Kwiecien, Piotr Brzoza, Pawel Majewski, Izabella Skulimowska, Kamil Bednarczyk, Joanna Cichy, Mateusz Kwitniewski

## Abstract

Chemerin is a chemoattractant protein with adipokine properties encoded by the retinoic acid receptor responder 2 (*RARRES2*) gene. It has gained more attention over the past few years due to its multilevel impact on metabolism and immune responses. The pleiotropic actions of chemerin include chemotaxis of dendritic cells, macrophages and natural killers (NK) subsets, bactericidal activity as well as regulation of adipogenesis and glucose metabolism. Therefore, reflecting the pleiotropic actions of chemerin, expression of *RARRES2* is regulated by a variety of inflammatory and metabolic mediators. However, for most cell types, the molecular mechanisms controlling constitutive and regulated chemerin expression are poorly characterized. Here we show that *RARRES2* mRNA levels in murine adipocytes are upregulated *in vitro* and *in vivo* by acute-phase cytokines, IL-1β and OSM. In contrast to adipocytes, these cytokines exerted a weak, if any, response in mouse hepatocytes, suggesting that the effect of IL-1β and OSM on chemerin expression is specific to fat tissue. Moreover, we show that DNA methylation controls the constitutive expression of chemerin. Bisulfite sequencing analysis showed low methylation levels within −735 to +258 bp of the murine *RARRES2* gene promoter in unstimulated adipocytes and hepatocytes. In contrast to these cells, the *RARRES2* promoter is highly methylated in B lymphocytes, cells that do not produce chemerin. Together, our findings reveal previously uncharacterized mediators and mechanisms controlling chemerin expression in various cells.

## Introduction

Chemerin is a small (18 kDa) multifunctional protein regulating different biological processes, including immune cell migration, adipogenesis, osteoblastogenesis, angiogenesis, myogenesis, and glucose homeostasis (1). Moreover, it shows broad-spectrum antimicrobial activity in human and mouse epidermis suggesting its role in maintaining skin barrier homeostasis (2, 3). Chemerin exerts most of these functions through the chemokine-like receptor 1 (CMKLR1) expressed by many cells including plasmacytoid dendritic cells (pDCs), macrophages, natural killers (NK) cells, adipocytes, hepatocytes, and keratinocytes (2, 4–9). Chemerin is secreted as pro-chemerin and circulates in plasma as an inactive precursor protein (Chem163S) that can subsequently be activated through posttranslational carboxyl-terminal processing by a variety of proteinases (10, 11).

The gene encoding chemerin is known as retinoic acid receptor responder 2 (*RARRES2*) (12), or tazarotene induced gene 2 (*TIG2*) since it was first discovered in tazarotene-treated psoriatic skin lesions (13, 14). Liver and adipose tissue are reported as the major sites of chemerin production; nonetheless, *RARRES2* mRNA is detected in many other tissues, including the adrenal gland, ovary, pancreas, lung, kidney, and skin (2, 15). Thus, reflecting the pleiotropic actions of chemerin, its association with inflammatory, metabolic and cell differentiation processes, and *RARRES2* tissue mRNA expression pattern, chemerin production is regulated by a variety of inflammatory and metabolic mediators that can be broadly classified as A) agonists of nuclear receptors (retinoids, vitamin D, glucocorticoids), B) factors mainly associated with metabolic processes (e.g. fatty acids, insulin, glucose) and C) immunomodulatory mediators (e.g. cytokines of acute or chronic inflammation and lipopolysaccharide (LPS) (1).

Although more than 20 years have passed since the *RARRES2* was discovered, still little is known about the molecular mechanisms regulating the transcription, translation, and secretion of chemerin in different cells. Chemerin expression may be constitutive or regulated (1). It is likely that these pathways are controlled differentially. For instance, adipocytes and hepatocytes show high constitutive *RARRES2* mRNA levels (15). However, pro-inflammatory cytokines, like TNFα, upregulate *RARRES2* mRNA mainly in adipocytes but not in hepatocytes (16). Eukaryotic gene expression can be regulated by a variety of mechanisms. This includes modifications of DNA (e.g. DNA methylation), binding of transcription factors to a gene promoter, alternative splicing, miRNAs and many others. Comprehensive functional analysis of the chemerin promoter has not yet been performed. Only two functional, *in silico* predicted, transcription factor binding sites have been validated *in vitro*. These studies identified functional response elements for the peroxisome proliferator-activated receptor γ (PPARγ), farnesoid X receptor (FXR), and sterol regulatory element-binding protein 2 (SREBP2) in the mouse chemerin promoter (17–19). These factors are regulated by lipids (PPARγ), bile acids (FXR) or free fatty acids (SREBP2). So far, the DNA methylation status of the *RARRES2* promoter and the role of methylation in the regulation of chemerin transcription has not yet been established.

Therefore, in this work we use adipocytes and hepatocytes to investigate constitutive and regulated expression of *RARRES2* after stimulation with interleukin 1β (IL-1β) and oncostatin M (OSM) pro-inflammatory cytokines. IL-1β and OSM are important acute-phase mediators that act on distant tissues to induce systemic effects of the acute-phase response to inflammatory stimuli like injury or infection. *In vitro*, IL-1β and OSM were shown to induce *RARRES2* mRNA and chemerin protein expression in cultured human skin equivalents (2) or 3T3-L1 adipocytes (IL-1β solely)(20). However, the effects of these cytokines on chemerin expression in primary hepatocytes and adipocytes have not been studied yet.

Here we show that acute-phase cytokines, IL1β and OSM, regulate chemerin expression in visceral white adipose tissue (vWAT) derived mouse primary adipocytes but not in primary hepatocytes, both *in vitro* and *in vivo*. Moreover, for the first time, we provide mechanistic information on the role of methylation in controlling constitutive chemerin expression.

## Experimental procedures

### Materials

If not stated differently, all chemicals were purchased form Sigma-Aldrich. RPMI-1640 medium was obtained from Biowest. DMEM, DMEM:F12 medium and PBS buffer were purchased from PAN Biotech. FBS was purchased from Gibco. BSA, EDTA and Trypan Blue were purchased from BioShop. Collagenase D was obtained from Roche, Germany. Mouse recombinant IL-1β and OSM were purchased from R&D Systems. Fc block (rat-anti-mouse CD16/32, #101310), biotin-conjugated rat-anti mouse CD45 (#103104), biotin-conjugated rat-anti mouse CD31 (#102404), and PE-conjugated rat-anti mouse Ly-6A/E (#108108) antibodies were purchased from Biolegend. LS columns, LD columns, anti-biotin and anti-PE magnetic beads were purchased from Miltenyi Biotec. PCR primers were obtained from Genomed, Poland.

### Animal studies

Male 8 - 12 week old C57BL/6 mice were used in these studies. Mice were housed under pathogen-free conditions in the animal facility at the Faculty of Biochemistry, Biophysics and Biotechnology of Jagiellonian University. IL-1β and OSM were injected intraperitoneally at a dose of 10 μg/kg BW and 160 μg/kg BW, respectively. After 48 h, the liver and epididymal white adipose tissue (eWAT) were isolated and subjected to RT-QPCR analysis. All experimental procedures were approved by the First Local Ethical Committee on Animal Testing at the Jagiellonian University in Krakow, Poland (permit no: 41/2014), in accordance with the Guidelines for Animal Care and Treatment of the European Community. Mice were sacrificed by overdose of anaesthesia (a mixture of ketamine and xylasine) followed by cervical dislocation.

### Primary hepatocytes isolation and culture

Primary hepatocytes were isolated from C57BL6 mice with a modified two-step perfusion method according to the protocol described by Seglen (21). Briefly, the mice were anesthetized with ketamine (100 mg/kg) and xylazine (10 mg/kg) i.p., and the abdomen was opened under sterile conditions. After cannulation of the portal vein, the liver was perfused with the solution I (100 μM EGTA in Krebs-Ringer (K-R) buffer), followed by the solution II (1 mg/ml collagenase D (Roche) in K-R buffer supplemented with 150 μM CaCl_2_). Then the liver was dissected, passed through a 100 μm cell strainer and centrifuged (60 g, 5min, 10°C). The isolated hepatocytes were suspended in DMEM:F12 medium supplemented with 10% FBS, 50 μg/ml gentamycin, 6 ng/ml insulin, and 400 ng/ml dexamethasone. Viable cells were counted using trypan blue staining and seeded on a collagen-coated plate at a density of 1 x 10^5^ cells/cm^2^. Cells were cultured at 37°C in presence of 5% CO_2_ for 4 hours. Afterwards, plated cells were washed with DMEM:F12 medium and stimulated with IL-1β (10 ng/ml) and OSM (50 ng/ml) for 48 hours. At the time of harvest, cell culture media supernatants were collected and RNA lysis buffer (Fenozol Plus, A&A Biotechnology, Poland) or RIPA buffer containing protease inhibitors (Roche) were added to the specified wells. Collected samples were subjected to RT-QPCR or ELISA analysis.

### Isolation and culture of adipose tissue-derived stromal vascular fraction (SVF)

Epididymal white adipose tissue depots were isolated from 8 - 10 week old male C57BL6 mice, minced and digested with collagenase D (3,5 mg/ml) in K-R buffer supplemented with 2% BSA and 150 μM CaCl2 for 1 hour at 37°C water bath with shaking every 5 min. Digested tissue were then centrifuged (280 g, 10 min, 15°C), washed, filtered through 100 μm cell strainer and centrifuged. Then the pellet was suspended in red blood cell lysis buffer (155 mM NH_4_Cl, 12 mM NaHCO_3_, 0,1 mM EDTA) for 3min at room temperature (RT), centrifuged and suspended in DMEM:F12 medium (20 % FBS, gentamycin 50 μg/ml). Isolated SVF cells were then seeded at a density of 9 x 10^4^ cells/cm^2^ in culture plate. Attached cells were washed and replenished with fresh medium after 24 hours to discard unattached dead cells or immune cells, and by day 3 more than 90% cells displayed typical fibroblastic morphology. Then, the SVF culture were subjected to adipocyte differentiation.

### Isolation and culture of adipogenic progenitors

Adipogenic progenitors (APs) were isolated from C57BL6 mice according to the protocol described by Lee at al. (22). Epididymal WAT was digested, filtered, and washed as described above. Adipocyte progenitors were then isolated using first negative selection of CD45 and CD31 followed by positive selection for SCA-1. Briefly, SVF cells were suspended in MACS buffer (0,5% BSA, 2mM EDTA, 50μg/ml gentamycin) and preincubated with Fc block (rat-anti-mouse CD16/32 mAbs, 45 μg/ml) followed by incubation with biotin-conjugated antibodies against CD45 (rat-anti-mouse CD45 mAb) and CD31 (rat-anti-mouse CD31 mAb), both 7,5 μg/ml. Then SVF cells were incubated with streptavidin-conjugated magnetic beads, washed and passed over a lineage depletion (LD) column to exclude endothelial cells and leukocytes. The flow-through was collected, washed, and incubated with PE-conjugated anti-SCA-1 (rat-anti-mouse Ly-6A/E mAb, 4,8 μg/ml) followed by anti-PE-conjugated magnetic beads. Cells were then washed and passed over a lineage selection (LS) column. Labeled cells were collected, suspended in DMEM:F12 medium (20 *%* FBS, gentamycin 50 μg/ml), counted, seeded on a cell culture flask and expanded. Then the cells were replated at a density of 2,5 x 10^4^ cells/cm^2^ and subjected to adipocyte differentiation. in adipocyte maintenance medium

### Adipocyte differentiation

Differentiation was induced when SVF cells or APs reached 90 % confluence. Cells were switched to adipocyte differentiation medium (DMEM:F12, 8% FBS, 8 μg/ml biotin, 50 μg/ml gentamycin, 1,15 μg/ml insulin, 80 μg/ml IBMX, 1,5 μg/ml troglitasone, 0,4 μg/ml dexamethasone) for 4 days. Subsequently, adipocyte differentiation medium was changed to adipocyte maintenance medium (DMEM:F12, 8% FBS, 8 μg/ml biotin, 50 μg/ml gentamycin, 1,15 μg/ml insulin) for another 7 days. The cells were stimulated with cytokines as described above.

### Isolation of B lymphocytes

Popliteal, axillary lymph nodes and spleen were dissected and pressed through the 40 μm mesh. Cells were washed with RPMI-1640, centrifuged (300 g, 6 min, 4°C), and pellet was suspended in red blood cell lysis buffer for 5 min at RT. Cells were centrifuged and suspended in MACS buffer (3 % FBS, 10 mM EDTA in PBS). B cell magnetic sorting was performed using MagniSort Negative Selection Protocol II (MagniSort Mouse B cell Enrichment Kit, Affymetrix). Negatively selected cells were suspended in PBS and DNA extraction was performed.

### 3T3-L1 cell line culture and cytokine stimulation

Primary mouse preadipocyte cell line 3T3-L1 was purchased from ATCC. 3T3-L1 cells were grown in DMEM medium supplemented with 10% FBS and gentamycin (50 μg/ml). Cells were seeded at a density of 8 x 10^3^ cells/cm^2^ in culture plate. After 24-hours medium was changed and cells were stimulated with IL-1β (10 ng/ml) and OSM (50 ng/ml) for 48 hours. At the time of harvest, cell culture media was removed and RNA lysis buffer was added. Collected samples were subjected to RT-QPCR analysis.

### RT-QPCR

Total RNA was extracted with Total RNA Zol-Out Kit (A&A Biotechnology, Poland) and converted to cDNA using NxGen M-MulV reverse transcriptase (Lucigen) with random primers (Promega). Real time PCR was performed on the CFX96 thermocycler (Bio-Rad, USA) using SYBR Green I containing universal PCR master mix (A&A Biotechnology) and primers specific for mouse: chemerin (5’-CTTCTCCCGTTTGGTTTGATTG, 5’-TACAGGTGGCTCTGGAGGAGTTC), SAA3 (5’- ACAGCCAAAGATGGGTCCAGTTCA, 5’- ATCGCTGATGACTTTAGCAGCCCA) and cyclophilin A (5’- AGCATACAGGTCCTGGCATCTTGT, 5’-CAAAGACCACATGCTTGCCATCCA). Relative gene expression normalized to cyclophilin A was calculated using the 2^−ΔΔCT^ method (23).

### ELISA

Chemerin levels in cell culture supernatants were quantified by mouse-specific ELISA. The MaxiSorp Nunc-Immuno Module (Thermo Fisher Scientific) strips were coated with rat-anti-mouse mAb (MAB23251, R&D Systems) in Tris-buffered saline (50 mM Tris-HCl pH 9.5, 150 mM NaCl). The plates were then washed with PBS containing 0.1% Tween 20, and nonspecific protein-binding sites were blocked with 3% BSA in PBS. Mouse recombinant chemerin was used as a standard. Chemerin was detected using biotin-conjugated rat-anti mouse chemerin mAb (BAM2325) followed by streptavidin-HRP (BD Science). The reaction was developed with TMB substrate (BD Science). The results were normalized to total protein content in the corresponding cell RIPA lysates (Quick Start Bradford Protein Assay, Bio-Rad). The ELISA detects both the 163S and 157S chemerin.

### Bisulfite genomic DNA sequencing

Genomic DNA was extracted from B cells, hepatocytes or differentiated APs using the GeneJET Genomic DNA Purification Kit (Thermo Fisher Scientific). Genomic DNA aliquots were then treated with sodium bisulfite using the EZ DNA Methylation-Direct Kit (Zymo Research). The targeted region of the *RARRES2* promoter was amplified with PCR (ZymoTaq PreMix, Zymo Research) using the following sets of primers specific to converted DNA: range −717/+229 (5’-GAGAGATTGAGTTGGGGAAATGAG-3’ sense, 5’-CCCCAACCTCTTTCTAATACCTTA-3’ antisense, 62,0°C), range −246/+154 (5’-ATGATAAAGGAAAGGTAAAGGAAAGATTGGG-3’ sense, 5’-AAACAACTCCCTAACAATTATTCCCTCTCACC-3’ antisense, 53,0°C), range −459/−160 (5’-GATGTTTGGTAGGTAGATGAAGGTAGTAGTTAGT-3’ sense, 5’-AACTACCATCAAAACAACTATCCCCAAC-3’ antisense, 58,9°C), range −813/−337 (5’-TAGGGAAAAGGTTTATTTGGTTAGTAGAGA-3’ sense, 5’-AAAAAAACTAAAACTCCTTCAATACCAAAA-3’ antisense, 50,2°C). PCR products were then separated on 2% agarose gel and extracted with Gel-Out Concentrator (A&A Biotechnology). Purified DNA were cloned into pTZ57R/T vector (InsTAclone PCR Cloning Kit, Thermo Fisher Scientific). After transformation and culturing of the competent bacteria (Top10 *E.coli*) overnight on an LB/agar/ampicillin plate, colonies (at least 8) were randomly selected, plasmids was recovered with GeneJET Plasmid Miniprep Kit (Thermo Fisher Scientific) and the DNA was sequenced with M13 common sequencing primers. The results were analyzed using QUMA online tool (RIKEN, Japan).

### Plasmid construction and *in vitro* methylation

The liver was dissected from C57BL6 mice and genomic DNA was isolated using the GeneJET Genomic DNA Purification Kit (Thermo Fisher Scientific). The *RARRES2* promoter was amplified using Phusion high-fidelity DNA polymerase (Thermo Fisher Scientific) and the following overlap extension PCR (OE-PCR) primers: 5’-CTCGAGGATATCAAGATCTGGCCTCGAAGCTTTCAGCTCCTCAGACAGGAA-3’ and 5’-GCTTTACCAACAGTACCGGATTGCCAAGTGGTACCTTGAAAATGATCAGGTTTGTT-3’ (−735 / +258; Chemerin Full). The resulting PCR product was subcloned into the promoterless pNL1.1[Nluc] vector (Promega, # N1001) using OE-PCR as described by Bryksin and Matsumura (24). Two additional constructs were then created using pNL1.1[Nluc]_Chemerin_Full as template and the following OE-PCR primers: 5’-CTCGAGGATATCAAGATCTGGCCTCGATCTGTCAAAAAACGGCTCCCTCAAGTG-3’ and 5’-CACTTGAGGGAGCCGTTTTTTGACAGATCGAGGCCAGATCTTGATATCCTCGAG-3’ (−252 / +258; Chemerin_Proximal), 5’-GAAGATCACCTGGTCAAGCGGGGCTTGGCAATCCGGTACTGTTGGTAAAGC-3’ and 5’-GCTTTACCAACAGTACCGGATTGCCAAGCCCCGCTTGACCAGGTGATCTTC-3’ (−735 / −253; Chemerin_Distal). The integrity and orientation of the inserts were confirmed by sequencing. Plasmids were amplified in *E.coli* and purified using Plasmid MIDI AX kit (A&A Biotechnology). For *in vitro* methylation studies the constructs were digested with NcoI and BglII (New England BioLabs Inc.), inserts and vectors bands were separated and extracted from the agarose gel. Inserts were then incubated for 1 hour at 37°C in the presence or absence of SssI methylase (New England BioLabs). The efficiency of the methylation reaction was verified by resistance to cleavage by the methylation-sensitive restriction enzyme HpaII (New England BioLabs). The methylated and mock-methylated chemerin promoter fragments were re-ligated into the parent vector, purified using Clean-Up Concentrator (A&A Biotechnology) and used for transfection.

### Transient transfection and luciferase assay

3T3-L1 cells were seeded at a density of 1,5 x 10^4^ cells in 24-well culture plate and 24 hours later cells were transfected using ViaFect transfection reagent (Promega). The total amount of DNA used for transfection per well was 0,8 μg, including approx. 0,5 μg of pNL1.1[Nluc] (equimolar concentrations of empty pNL1.1[Nluc], pNL1.1[Nluc]_Chemerin_Full, pNL1.1[Nluc]_Chemerin_Proximal or pNL1.1[Nluc]_Chemerin_Distal) and 0,3 μg of the pGL4.54 [luc2/TK] vector (Promega) expressing the Firefly luciferase that was used as an internal control. For methylation studies, cells were transfected with 0,5 μg of ligation reactions from the methylated/mock-methylated pNL1.1[Nluc]/ chemerin promoter constructs and 0,3 μg of the pGL4.54 [luc2/TK] vector. The amount of DNA per well was equalized using mock plasmid DNA (pcDNA3.1, Promega). Forty-eight hours later, the cells were harvested by scraping into a passive lysis buffer (Promega). NanoLuc and Firefly activities in cell lysates were measured using Nano-Glo Dual-Luciferase Reporter Assay System (Promega) according to the manufacturer’s protocols.

### Statistical analysis

All the data was analyzed using STATISTICA 13, visualized with Prism GraphPad and present as mean ± SD or mean ± SEM. Student’s t-test was used for comparison between two groups. For multiple comparisons either ANOVA with Bonferroni post-hoc test or Kruskal-Wallis ANOVA was used. For experiments with repeated measures over time, ANCOVA with Bonferroni post-hoc test was used instead. Differences were considered statistically significant for p-value <0,05 and marked with an asterisk.

## Results

### IL-1β and OSM stimulation upregulates *RARRES2* expression in murine adipocytes but not hepatocytes

White adipose tissue and the liver have been confirmed by multiple studies as key sites of chemerin production (1). Both organs contribute to fatty acid metabolism, respond to numerous cytokines and are involved in the pathogenesis and pathophysiology of obesity (25) which is linked to elevated systemic levels of chemerin (26). Among potential regulators of systemic chemerin levels are acute-phase mediators OSM and IL-1β which, as we showed previously, regulate chemerin levels in human epidermis-like cultures (2). Therefore, we determined if IL-1β and OSM affect chemerin expression in adipocytes and hepatocytes *in vitro* and *in vivo*. Treatment of vWAT-derived mouse adipocytes with OSM and/or IL-1β resulted in statistically significant and comparable upregulation of *RARRES2* mRNA by each stimulus (Fig.1A), as well as a trend to higher chemerin protein levels (Fig. 1B) after 48-hour stimulation. In contrast to adipocytes, downregulation of *RARRES2* mRNA was detected in hepatocytes in response to OSM or OSM + IL-1β but not IL-1β alone. Cytokines did not affect chemerin protein levels in hepatocytes cell culture supernatants (Fig. 1B). Similar results were obtained after *in vivo* IL-1β + OSM administration. *RARRES2* mRNA was upregulated only in vWAT (Fig. 1D–E) but not in the liver. We confirmed that primary hepatocytes and mouse liver responded to stimulation since *SAA3* mRNA, encoding an acute-phase protein, was markedly elevated (Fig. 1C and 1F). Together, these results suggest that the effect of the cytokines that results in upregulation of chemerin expression is specific to fat tissue.

**Figure 1.**
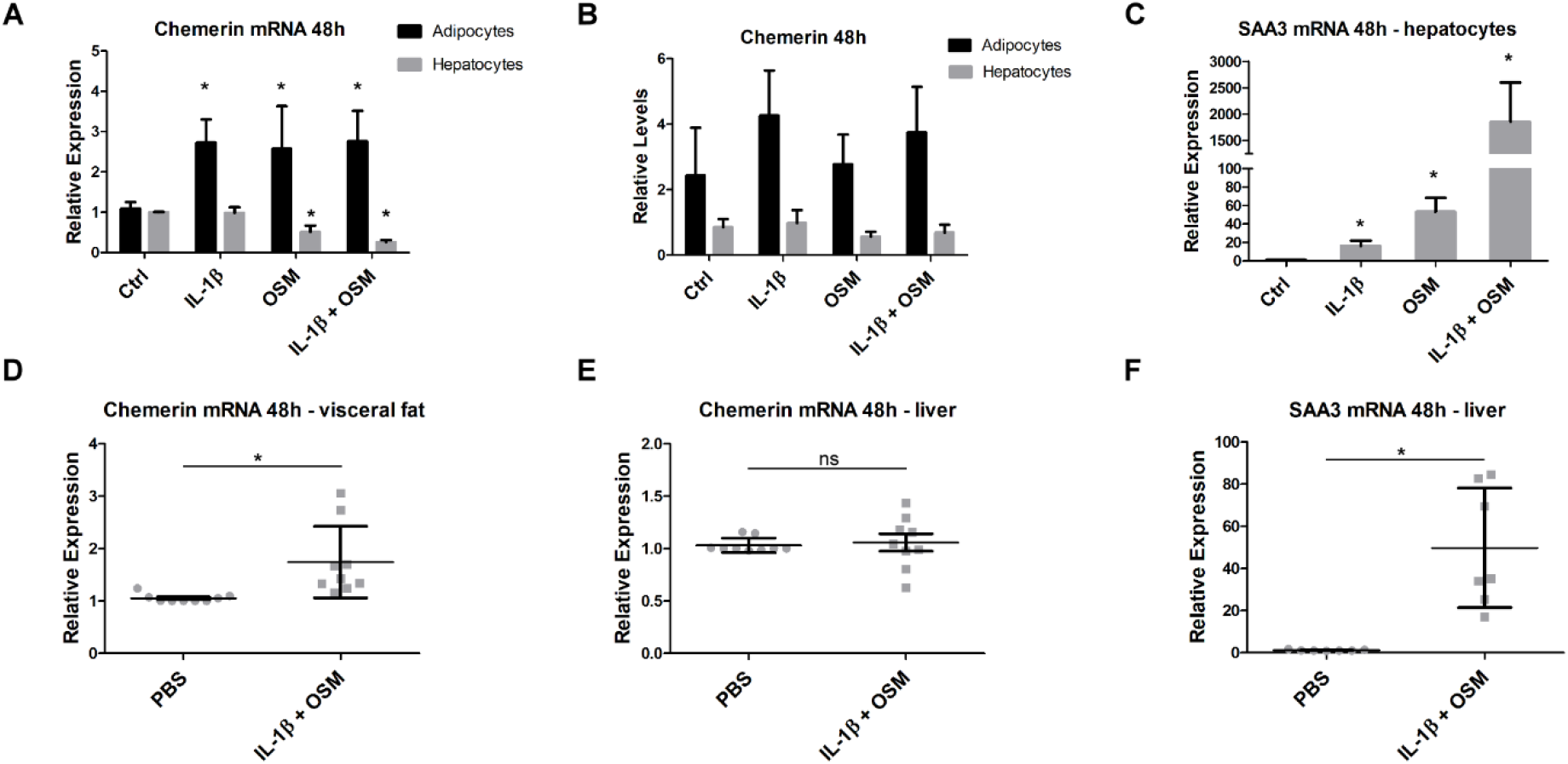
Acute-phase cytokines upregulate chemerin expression in adipocytes of white adipose tissue but not in hepatocytes. Primary mouse hepatocytes or the stromal vascular fraction (SVF) of visceral white adipose tissue (vWAT) were isolated. SVF cells were differentiated to obtain a mature adipocyte cell culture. Then, the cells were treated with IL-1β (10 ng/ml), OSM (50 ng/ml) or a combination for 48-hours. The levels of chemerin (A) and SAA3 (C) mRNA were determined using RT-QPCR. Relative expression of stimulated cells over the control is shown. Levels of secreted chemerin were determined in parallel in conditioned media by ELISA (B). Data are presented as the mean ±SD. Statistical significance between the control and treated cells is shown by an asterisk; *p<0.05 by ANOVA followed by a Bonferroni post hoc test. *In vivo*, IL-1β and OSM were injected intraperitoneally at a dose of 10 μg/kg BW and 160 μg/kg BW, respectively. After 48 h, the liver and visceral white adipose tissue (vWAT) were isolated and subjected to RT-QPCR analysis. The levels of chemerin mRNA in vWAT (D) or liver (E) and SAA3 (F) were determined. Data are presented as the mean ±SD. Statistical significance between the control (PBS) and the cytokine treated animals is shown by an asterisk; *p<0.05 by the two-tailed Student’s t test. All the experiments were repeated at least three times.

### Adipocytes are key cells expressing chemerin after stimulation with acute-phase cytokines

The stromal vascular fraction (SVF) of adipose tissue consists of a heterogeneous population of cells that includes adipocyte precursors, hematopoietic stem cells, endothelial cells, fibroblasts, and immune cells (27). Therefore, we next asked if adipocytes are the main cells that express chemerin in response to acute-phase cytokines. Indeed, we found a similar chemerin expression pattern and levels in both vWAT-derived adipocytes and in adipocytes derived from sorted SCA-1^+^ adipogenic progenitors (Fig. 2A–D). Chemerin transcript levels continue to rise throughout a 5-day time course. Stimulation with IL-1β and IL-1β + OSM showed the highest mRNA levels (approx. 12–13-fold increase over the 1-day control) in both types of adipocyte cell culture (Fig. 2A and 2C). However, statistically significant upregulation of chemerin protein levels was observed only in IL-1β + OSM treated vWAT-derived adipocytes or in the cell culture of sorted adipogenic precursors after IL-1β and IL-1β + OSM stimulation. Accordingly, we concluded that IL-1β and OSM regulate chemerin expression primarily in adipocytes. Chemerin protein production was the highest in response to IL-1β + OSM, suggesting additive effects.

**Figure 2.**
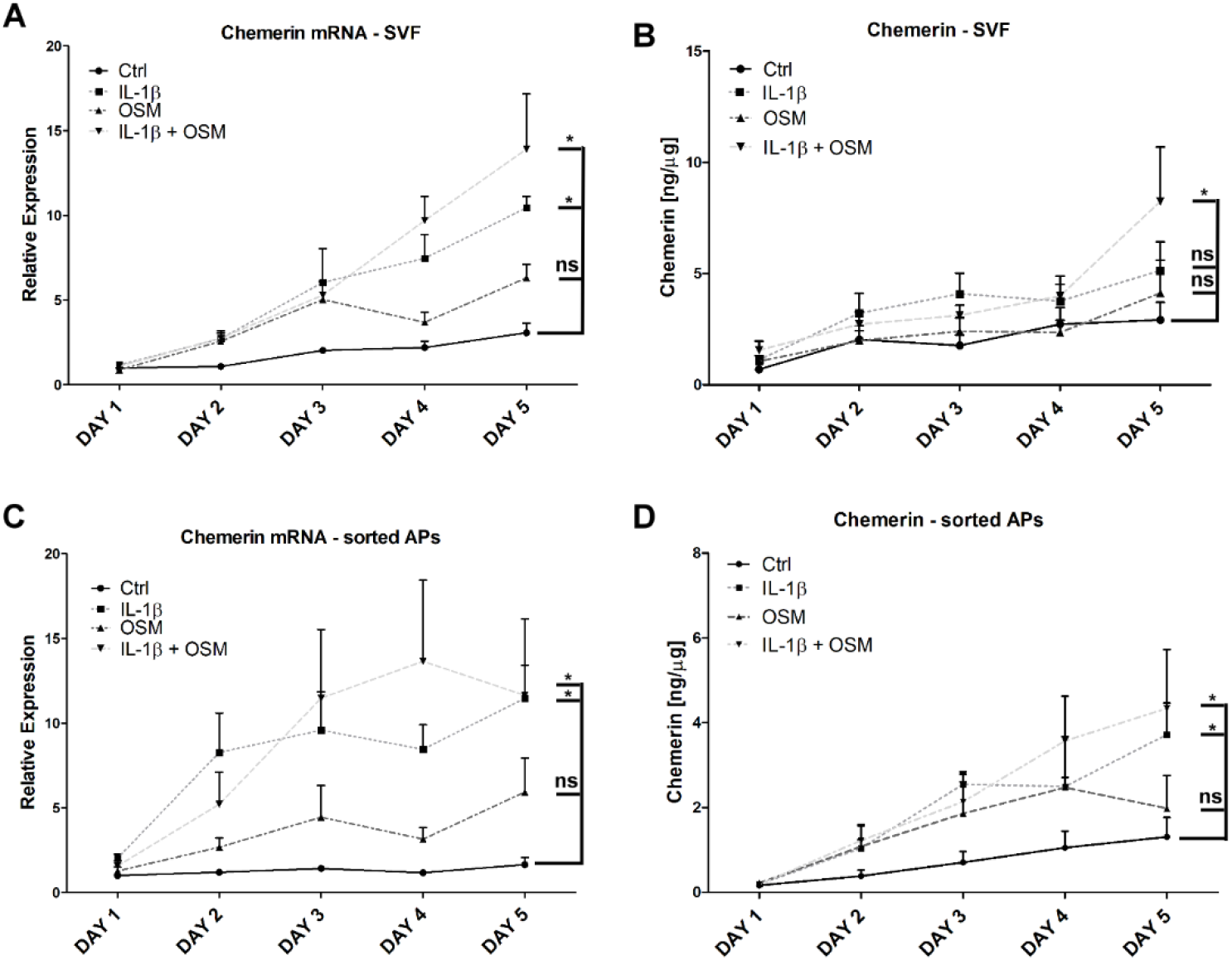
Adipocytes are key cells expressing chemerin after stimulation with IL-1β + OSM. vWAT or SCA-1+ adipogenic progenitors (AP) were isolated. The cells were differentiated to obtain a mature adipocyte cell culture. Then, the cells were treated with IL-1β (10 ng/ml), OSM (50 ng/ml) or a combination for up to 5 days. The levels of chemerin mRNA (A, C) were determined using RT-QPCR. Relative expression of stimulated cells over the control is shown. Levels of secreted chemerin were determined in parallel in conditioned media by ELISA (B, D). Data are presented as the mean ±SEM. Statistical significance between the control and the treated cells is shown by an asterisk; *p<0.05 by ANCOVA followed by a Bonferroni post hoc test.

### DNA methylation affects *RARRES2* expression

The functional significance of DNA methylation in the regulation of *RARRES2* transcription has not yet been elucidated. Therefore, using bisulfite sequencing we determined the methylation status of the murine *RARRES2* gene in unstimulated and cytokine-treated adipocytes and hepatocytes. B cells were used as a reference since these cells do not produce chemerin. We found 17 CpG sites located within −735/+258 bp of *RARRES2* (Fig. 3A). However, computational analysis of this sequence did not identify any CpG island (data not shown). The results showed low methylation levels within the murine chemerin promoter in unstimulated adipocytes and hepatocytes (Fig. 3B). In contrast to these cells, the chemerin promoter is highly methylated in B lymphocytes. Interestingly, upregulation of chemerin expression after stimulation of adipocytes with IL-1β and OSM correlates with the statistically significant increase in the average methylation level of the *RARRES2* gene promoter within −649/−257 bp (Fig. 3C) but not −239/+222 bp (Fig. 3D). The methylation pattern of the chemerin promoter was not changed in stimulated primary cultures of mouse hepatocytes.

**Figure 3.**
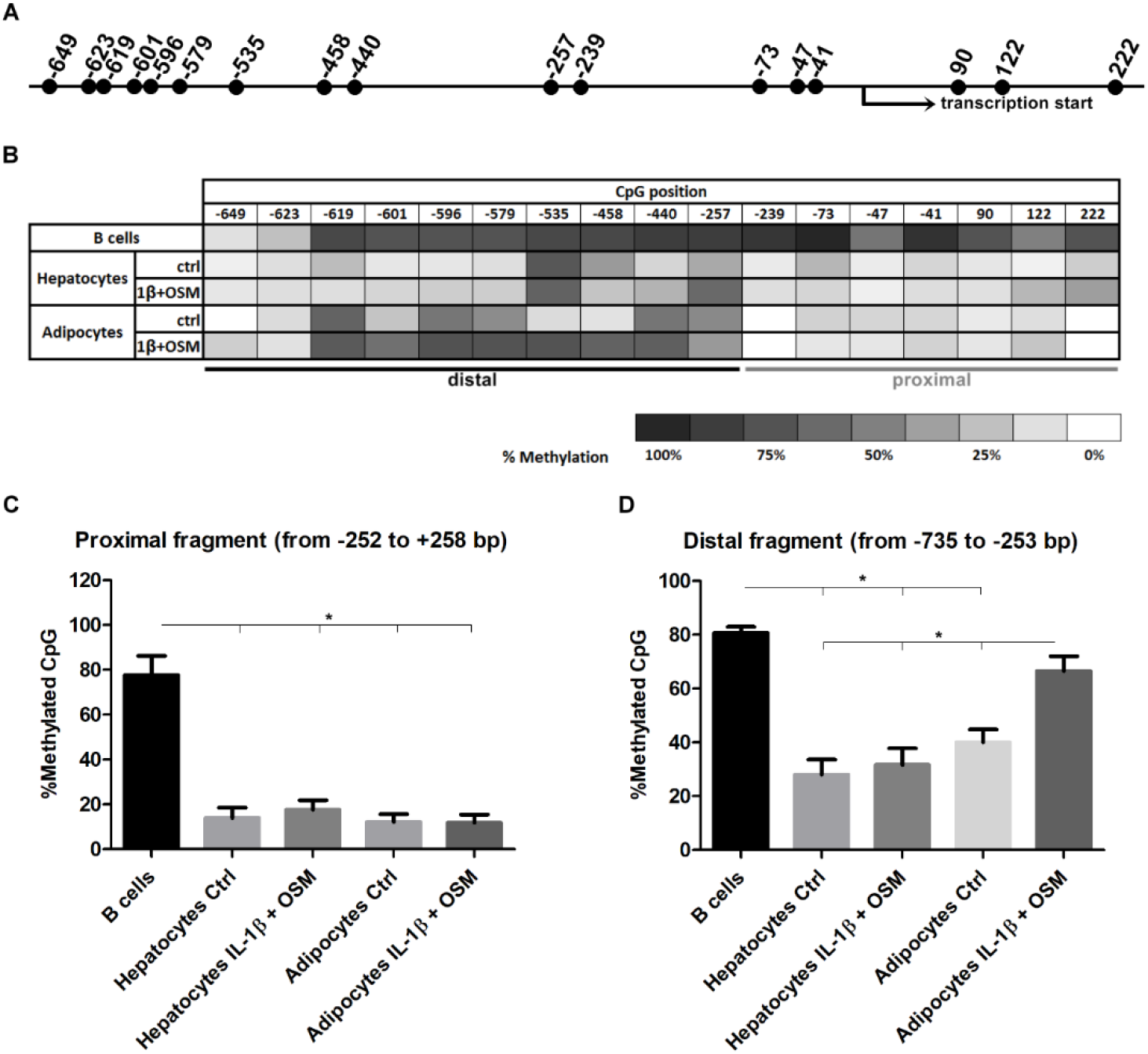
Chemerin transcription is linked to the DNA methylation status of the *RARRES2* promoter from −735 to +258 bp. Primary mouse hepatocytes and mature adipocytes derived from sorted APs were treated with IL-1β (10 ng/ml) and OSM (50 ng/ml) for 48 hours. B cells, that do not produce chemerin, were used as a control. The DNA methylation status of the *RARRES2* promoter was analyzed by bisulfite sequencing. The localization of CpG sites (A) and the DNA methylation levels of each CpG site from −735 to +258 bp of the *RARRES2* promoter are shown (B). The differences in the average methylation level of the proximal (C) and distal (D) parts of the promoter were analyzed using the nonparametric Kruskal-Wallis test; *p<0.05. Data are presented as the mean ±SEM.

### The proximal part of the *RARRES2* promoter is a key regulator of transcription

To test how cytokines or DNA methylation can regulate transcription of the *RARRES2* we made three reporter constructs containing different portions of chemerin promoter sequences: Chemerin_Full (position −735/+258 bp relative to the transcription start site), Chemerin_Proximal (position −252/+258 bp), and Chemerin_Distal (position −735/−253 bp) (Fig. 4A). We used 3T3-L1 adipocyte precursors for transient transfection since these cells produce chemerin and respond to cytokine stimulation like differentiated adipocytes (Fig. 4B). Chemerin_Proximal and Chemerin_Full had the highest promoter activity, as shown by luciferase assay (Fig. 4C). The Chemerin_Distal construct had similar promoter activity compared to the empty vector. We observed downregulation of promoter activity of Chemerin_Proximal and Chemerin_Full after 48-hour stimulation of 3T3-L1 cells with IL-1β + OSM. To investigate the role of DNA methylation in the regulation of chemerin expression, we excised the promoter sequence from pNL1.1[Nluc] constructs with restriction enzymes, treated the eluted DNA with SssI methylase or left it untreated, and religated the methylated and unmethylated DNA inserts into the parent vector. Promoter activity of the methylated Chemerin_Proximal was five-times lower than that of the unmethylated construct (Fig. 4D). In line with the results of bisulfite sequencing (Fig. 3), promoter activity of the methylated Chemerin_Distal was almost four-times higher than the unmethylated construct. Together, these data suggest that the DNA sequence of the *RARRES2* promoter located from −252 to +258 bp is critical for chemerin transcription and its methylation suppresses the transcription. Interestingly, the distal part of the chemerin promoter (position −735/−253 bp) shows minimal activity but seems to play a regulatory role since DNA methylation increases its activity.

**Figure 4.**
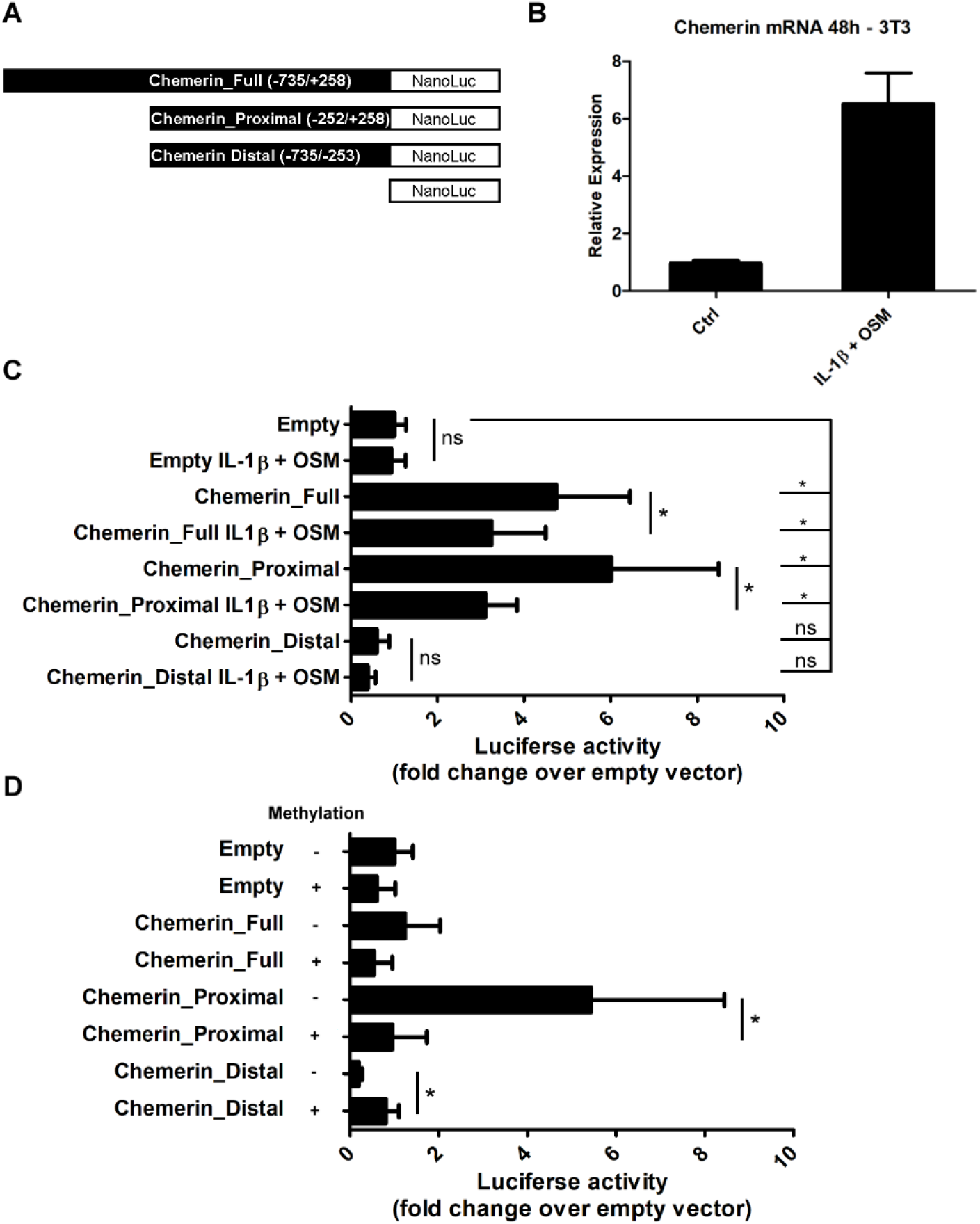
The proximal part of the chemerin promoter is a key regulator of transcription. The murine *RARRES2* promoter regions (from −735 to +258 bp) were cloned into the pNL1.1[Nluc]-Basic vector (A). 3T3-L1 adipocyte precursors were used for transient transfections since these cells respond to IL-1β + OSM stimulation similarly to the primary adipocytes (B). 3T3-L1 cells were transfected with different chemerin promoter constructs (C, D). The results are expressed as the fold-change in relative luciferase units (RLUs) relative to the empty vector. Statistical significance between the empty vector and the promoter construct is shown by an asterisk; *p<0.05 by ANOVA followed by a Bonferroni post hoc test. The Mann–Whitney test was used to analyze the statistical differences between the control and IL-1b + OSM treated cells or methylated/mock methylated vectors. Data are presented as the mean ±SD.

## Discussion

In this study, we demonstrate differential *RARRES2* gene regulation in IL-1β and OSM treated murine adipocytes and hepatocytes, both *in vitro* and *in vivo*. Our data also reveal the key importance of methylation of specific regions of the chemerin promoter in the regulation of chemerin expression. For the first time we show that cells expressing chemerin have a different methylation profile of *RARRES2* compared to cells that do not produce this protein.

Pro-inflammatory cytokines, including TNFα, INFγ, IL-6, IL-1β, and OSM, are essential mediators of adaptive and innate immune responses (28, 29). Elevated serum levels of these cytokines are linked to psoriasis (30), rheumatoid arthritis (31), cancer (32), and obesity (33), pathological conditions with postulated chemerin involvement (6, 26, 34, 35). Therefore, elucidating the underlying mechanisms involved in the regulation of chemerin expression is of considerable interest.

Chemerin is expressed at the highest levels in the liver and white adipose tissue (1), organs that contribute to fatty acid metabolism, respond to numerous cytokines, and are involved in the pathophysiology of obesity (25). Many studies confirmed a relationship between serum chemerin and fat mass in overweight/obese individuals and correlations with obesity-associated traits, like low-grade inflammation, hypertension, and insulin resistance in some but not all of the patient cohorts studied (36). Both cytokines, IL-1β and OSM, are linked with low-grade inflammation in obesity. IL-1β can influence the metabolic function of adipocytes and induce insulin resistance (37). OSM inhibits adipocyte development, modulates glucose and lipid metabolism (38), and contributes to hepatic insulin resistance and the development of non-alcoholic steatohepatitis (NASH) (39). IL-1β and OSM can induce production of C-reactive protein (CRP) in the liver, independently of IL-6 (25). *In vitro*, IL-1β increased chemerin protein concentrations in conditioned media from 3T3-L1 adipocytes and the expression was upregulated in these cells in a time- and dose-dependent manner (20). IL-1β and OSM upregulated chemerin expression in human skin cultures (2). Our studies revealed that a combination of both cytokines considerably increased *RARRES2* mRNA and protein levels in cell cultures of murine vWAT-derived adipocytes and differentiated SCA1+ APs but not in primary hepatocytes. *In vivo* experiments confirmed the results shown *in vitro. RARRES2* mRNA was elevated in vWAT but not in liver. vWAT plays a key metabolic and endocrine role in overweight individuals (40). This led us to the conclusion that adipocytes have an important role in sensing and responding to cytokines by supporting chemerin production. In contrast to adipocytes, IL-1β and OSM did not affect chemerin expression in hepatocytes. So far, only free fatty acids (FFA) and GW4064, a synthetic farnesoid X receptor (FXR) agonist, were shown to influence chemerin expression in hepatocytes (18). FXR is known as a primary bile acid nuclear receptor, which plays a critical role in maintaining lipid and glucose homeostasis (18). A pro-inflammatory cytokine, TNFα, was reported to affect chemerin expression in various cells like adipocytes (16), human synoviocytes (35), or fetal human intestinal cells (41). However, it did not induce the expression of chemerin in the liver or primary hepatocytes, which implicated adipocytes as a potential source of elevated serum chemerin after TNFα exposure *in vivo* (16). Taken together, the data indicate different regulation of chemerin expression in adipose tissue and the liver, and suggest that regulation of the expression of chemerin within adipose tissue may be of importance for the pathogenesis of obesity and other inflammatory diseases.

DNA methylation is an essential epigenetic mechanism involved in gene regulation. Epigenetic regulation by CpG methylation in the promoter region is often associated with both tissue-specific and heterogeneous expression of genes (42). DNA methylation can cause transcriptional silencing of genes by inhibiting the binding of transcription factors (TFs) to regulatory sequences or by the binding of methyl-CpG binding proteins. Methyl-CpG binding proteins bind to methylated DNA and recruit histone deacetylases that induce deacetylation of histones, leading to chromatin condensation and transcriptional repression (43). However, methylation in the immediate vicinity of the transcriptional start sites blocks initiation, but methylation in the gene body does not block and might stimulate transcription elongation (42). Even the methylation status of a single CpG locus can modulate gene expression (44, 45). Here we show, for the first time, that CpG sites located from −619 to +222 bp of the *RARRES2* promoter are highly methylated in B cells but not hepatocytes and adipocytes. This suggests that DNA methylation controls constitutive chemerin expression in cells of different origins. Surprisingly, elevated mRNA levels in IL-1β + OSM stimulated adipocytes, but not hepatocytes, correlated with increased methylation within CpG sites located from −619 to −257 bp of the *RARRES2* promoter. Therefore, we divided the chemerin promoter into two parts: proximal (−252/+258 bp) and distal (−735/−253 bp). Proximal and distal regions have different methylation levels and transcription factor binding sites. The average percentage methylation levels for the proximal region is less than 14 % in both unstimulated adipocytes and hepatocytes since the distal part of the promoter shows higher average methylation, varying from 28 % (hepatocytes) to 40 % (adipocytes). The proximal part of the murine chemerin gene promoter has two important functional transcription factor binding sites: the PPAR response element (PPRE) (19) and FXR response element (FXRE) (18). Notably, the location and sequence of the functional FXRE overlaps with the PPRE (1). PPRE is responsible for the regulation of chemerin gene expression by PPARγ, a master regulator of adipogenesis (19). FXR, expressed at high levels in the liver, is the main receptor for bile acids, natural detergents involved in the absorption of dietary fat and fat-soluble vitamins (1). The distal part of the *RARRES2* promoter has a functional SREBP2 binding site located close to the proximal part of the promoter (17). SREBP2 activates genes in cholesterol biosynthesis (17). Listed TF binding sites are strictly linked with factors regulating chemerin expression in adipocytes and hepatocytes. Indeed, luciferase assays showed that the proximal part of the chemerin promoter is a key regulator of chemerin expression. Methylation of the CpG sites located within this region diminished luciferase activity. The distal part of the chemerin promoter is poorly characterized, and did not show any important activity in luciferase assays. However, methylation of the Chemerin_Distal construct increased luciferase activity compared with the unmethylated vector which resembles the *in vitro* bisulfite sequencing results of Il-1β and OSM stimulated adipocytes.

Together, our findings reveal novel insights into the mechanisms and factors regulating chemerin expression and secretion. The results suggest that DNA methylation controls the constitutive expression of *RARRES2* in murine adipocytes and hepatocytes, and the proximal region of the gene promoter is the key regulator. Moreover, for the first time we show that acute-phase cytokines upregulate *RARRES2* mRNA in adipocytes but not hepatocytes, both *in vitro* and *in vivo*, indicating distinct transcriptional regulatory programs in these cells. Further studies are needed to elucidate the importance of the distal part of the *RARRES2* promoter since cytokines elevated the methylation level of these regions in murine adipocytes.

## Abbreviations

IL: interleukin
OSM: oncostatin M
SVF: stromal vascular fraction
vWAT: visceral white adipose tissue

## Author Contribution

K.K, J.C. and M.K. conceived and designed the experiments; K.K., P.B., P.M., I.S., K.B. and M.K. performed experiments; K.K., P.B., J.C. and M.K. analyzed data; M.K. wrote the manuscript. All authors have approved the manuscript.

## Acknowledgments

The Faculty of Biochemistry, Biophysics, and Biotechnology of the Jagiellonian University is a partner of the Leading National Research Center (KNOW) supported by the Polish Ministry of Science and Higher Education.

